# Simultaneous whole-cell recording and calcium imaging to reveal electrically coupled neurons in *Xenopus* tadpoles

**DOI:** 10.64898/2026.03.04.707658

**Authors:** Bella Xu Ying, Maarten F. Zwart, Wen-Chang Li

## Abstract

Neuronal populations connected by gap junctions can be revealed via dye coupling of small molecules like neurobiotin and lucifer yellow. However, the extent of dye diffusion between neurons varies with connexin subtype, loading method, and neuromodulation. Due to the increasing availability of GCaMP transgenic animals, we explore the possibility of revealing gap junctional coupling using Ca^2+^ imaging in the *Xenopus laevis* tadpole motor system. Reliable axo-axonal electrical coupling was previously found in excitatory descending interneurons (dINs) using paired recordings but not with neurobiotin dye coupling. Here, we made whole-cell patch-clamp recordings with Ca^2+^-supplemented intracellular solution to load Ca^2+^ into GCaMP6s-expressing neurons, followed by Ca^2+^ imaging to detect potential Ca^2+^ diffusion across coupled neurons. Successful membrane breakthroughs led to transient fluorescence increases in the patched neuron. However, increasing the Ca^2+^ concentration promoted membrane resealing and rapid loss of whole-cell recordings. Regardless of recording duration, loading-triggered fluorescence only lasted up to three minutes, suggesting rapid Ca^2+^ clearance. Pharmacologically blocking sarcoplasmic /endoplasmic reticulum Ca^2+^-ATPases and plasma membrane Na^+^/Ca^2+^ exchangers did not prolong fluorescence, although sustained fluorescence was achieved with positive current injections. Counter to our expectations, fluorescence increases in Ca^2+^-loaded dINs did not spread to neighboring dINs. Robust intracellular Ca^2+^ regulation mechanisms, membrane resealing, and long dIN axons likely hindered intercellular Ca^2+^ diffusion. Therefore, this approach is not appropriate for revealing electrical coupling within this system.

## Introduction

Gap junctions are essential in cell-to-cell communication. In non-neural tissue like epithelial cells, glia, and cardiac muscle cells, gap junctions are found ubiquitously and form homogeneous cell networks (Severs, 1994; Orthmann-Murphy, Abrams and Scherer, 2008; Giepmans and van IJzendoorn, 2009). In contrast, neuronal gap junctions, or electrical synapses, are mostly found among specific cell types (Söhl, Maxeiner and Willecke, 2005). These channels play important roles in activity synchronization within neural populations (Zhang *et al*., 2009), development (Cao, Liu and Yu, 2023), and cell survival (Belousov *et al*., 2017). Consequently, gap junction defects have been associated with various neurodegenerative diseases such as Alzheimer’s and Parkinson’s (Sánchez *et al*., 2020; Xing and Xu, 2021; Choudhury *et al*., 2022). However, the location and function of electrical synapses in the nervous system remain largely unknown. The different gap junction expression patterns across neuron types require reliable and effective methods to determine the extent of electrical coupling.

Standard methods for revealing gap junction-connected cells include paired recordings and dye coupling. In paired recordings of coupled cells, applying current injections to one cell results in trans-junctional voltage changes in the other (Bennett *et al*., 1963). Although highly sensitive, the technical challenges involved in performing paired recordings and its limitation to two cells per experiment reduce efficiency and throughput (Abbaci *et al*., 2008). In dye coupling experiments, small dyes or tracers like neurobiotin, biocytin, and lucifer yellow can reveal coupled cells or networks by diffusing across gap junctions (Stephan, Eitelmann and Zhou, 2021). This is less technically challenging than paired recordings, but the extent of dye coupling hinges on several uncertainties. First, dye microinjections or scrape loading can cause cell damage, making the amount of dye loaded difficult to standardize (Abbaci *et al*., 2008). Consequently, damage can trigger neuromodulation and homeostatic mechanisms that affect gap junction permeability (Ziambaras *et al*., 1998; Bloomfield and Völgyi, 2009). Moreover, gap junctions have different dye permeabilities depending on their connexin composition. For example, lucifer yellow can pass through Cx40, Cx43, and Cx45 gap junctions but not Cx30.2 (Rackauskas, Verselis and Bukauskas, 2007). These dependencies present challenges in studying electrical synapses, especially in cells with gap junctions of unknown connexin makeup.

In recent years, new genetically-encoded methods have emerged, many of which use a “generator-reporter” system (Dong, Liu and Li, 2018). Ion channels or pumps are expressed in “generator” cells and sensor proteins in coupled “reporter” cells. The experimenter can then trigger an inflow of ions into the “generator” (e.g., via optogenetics), and the ions’ diffusion across gap junctions is detected by the “reporter” (e.g., via fluorescence sensor proteins). These methods are feasible in *Drosophila melanogaster* (Wu *et al*., 2019) and cell cultures (Kang and Baker, 2016) where sophisticated genetic toolkits are available. However, for many model systems in which similar tools are lacking, such as *Xenopus laevis*, studying gap junctions is limited to conventional techniques like paired recordings and dye coupling. Previously, paired recordings from the swimming motor circuits in *Xenopus* tadpoles revealed electrical coupling amongst descending interneurons (dINs) through potential axo-axonal gap junctions (Li, Roberts and Soffe, 2009; Zhang *et al*., 2009). Reliable electrical coupling was found using paired recordings between 55 dIN pairs but neurobiotin dye coupling rarely diffused beyond the injected cell (Li, Roberts and Soffe, 2009). Due to inconsistencies across methods, our understanding of dIN coupling at the population level remains limited, hindering our functional understanding of their role in locomotion.

We attempted to explore an alternative method for revealing electrical coupling among *Xenopus* tadpole dINs: using whole-cell electrodes to load Ca^2+^ into GCaMP-expressing neurons and Ca^2+^ imaging to detect trans-junctional Ca^2+^ diffusion to potentially coupled cells. Ca^2+^ ions (40 Da) are smaller than neurobiotin (286 Da) (Mills and Massey, 1998) and should diffuse more readily through gap junctions. Moreover, we used a common pan-neuronal GCaMP6s line available to many model systems, removing the need for complex genetic tool development. We loaded Ca^2+^ into single cells and monitored changes in GCaMP6s fluorescence upon membrane breakthrough. First, we showed that loading Ca^2+^ produced a rapid fluorescence increase in the recorded neuron with an average halftime of 103 seconds. Raising the Ca^2+^ concentration and applying pharmacological blockers of Ca^2+^ pumps, which normally remove cytoplasmic Ca^2+^, did not prolong the fluorescent signal. Instead, loading high Ca^2+^ concentrations promoted resealing of the cell membrane within the electrode and rapid loss of whole-cell recordings. Although positive current injections could sustain fluorescence increases within the loaded cell, loading dINs with Ca^2+^ did not lead to detectable fluorescence increases in neighboring dINs. We therefore conclude that this is not a viable method for revealing electrical coupling between *Xenopus* tadpole dINs.

## Materials and Methods

### Animals

Pairs of adult *Xenopus laevis* frogs were induced to breed via injections of 1000 U/ml human chorionic gonadotropin hormone (HCG, Sigma-Aldrich, Gillingham, UK). Fertilized embryos were collected and raised at temperatures between 16-22 ºC to stagger developmental rates. Stage 37/38 tadpoles were used in all experiments. All procedures were approved by the University of St Andrews Animal Welfare Ethics Committee and comply with UK Home Office regulations (license number PP2896979).

### Dissection

Tadpoles were anaesthetized for 10-20 seconds in 0.1% 3-aminobenzoic acid ester (MS-222, Sigma-Aldrich, Gillingham, UK) to allow opening of the dorsal fin, then immobilized for 25 minutes with 13 µM α-bungarotoxin (Sigma-Aldrich, Gillingham, UK). Immobilized tadpoles were pinned to a rotatable Sylgard dissection dish filled with extracellular saline (in mM: 115 NaCl, 10 HEPES, 3 KCl, 2 CaCl_2_, 1 MgCl_2_, 2.4 NaHCO_3_; adjusted with 5M NaOH to pH 7.4, with an osmolarity of ∼257 Osm/L). Most of the yolk sac and the rostral trunk skin covering both sides of myotome blocks were dissected out. The skin covering the hindbrain was removed and a slit was made dorsally along the roof of the fourth ventricle and rostral spinal cord to open respective neurocoeles. Ependymal cells lining the neurocoele were removed with a fine dissection needle to expose hindbrain and spinal cord somas. The left side of the caudal hindbrain and rostral spinal cord was removed (hemicord) to improve visibility using the transmitted light on an E600FN microscope (Nikon, Tokyo, Japan).

### Whole-cell patch-clamp recordings and Ca^2+^ loading

After dissection, tadpoles were transferred to a rotatable recording dish and whole-cell patch-clamp recordings in current-clamp mode were made following established methods (Li, 2021). Electrodes were pulled with a P-87 pipette puller (Sutter Instrument, California, USA) with DC resistances between 10-20 MΩ. The standard intracellular solution contained (in mM): 100 K-gluconate, 2 MgCl_2_, 10 EGTA, 10 HEPES, 3 Na_2_ATP, 0.5 NaGTP, 3.1 neurobiotin; adjusted with KOH to pH 7.3, with an osmolarity of ∼239 Osm/L. For Ca^2+^ loading experiments, CaCl_2_ was added and Ca^2+^ chelators EGTA, K-gluconate and Na_2_ATP were replaced with K-CH_3_SO_3_ and MgATP (Woehler, Lin and Neher, 2014) for final concentrations of (in mM): 107 K-CH_3_SO_3_, 1 MgCl_2_, 2 CaCl_2_, 10 HEPES, 3 MgATP, 0.5 NaGTP, 3.1 neurobiotin; adjusted with KOH to pH 7.3, with an osmolarity of 243 Osm/L. In the 10 mM Ca^2+^ loading experiments, the intracellular solution with 10 mM CaCl_2_ had an osmolarity of 267 Osm/L. We therefore increased the extracellular saline osmolarity accordingly by raising NaCl to 127 mM for an osmolarity ∼281 Osm/L. Epifluorescence Ca^2+^ imaging was performed with an Andor Neo5.5 CMOS camera and Andor Solis software (Oxford Instruments, Oxfordshire, UK). Blue light was supplied via a 480nm LED array module on a pE-1 system (CoolLED, Andover, UK). LED lighting and image capture were synchronized through TTL pulses configured through Signal software (Cambridge Electronic Design, Cambridge, UK). Intracellular signals were amplified with a Multiclamp 700B amplifier (Molecular Devices, CA, USA). Fictive swimming was monitored by recording motor nerve activity from myotome clefts with a glass suction electrode, amplified with a 4-channel extracellular amplifier (AM Systems, WA, USA), and digitized with a Power1401 Digitizer (Cambridge Electronic Design, Cambridge, UK). All electrophysiological recordings were acquired at a 10 kHz sampling rate using Signal.

The seal between the patch-clamp electrode and neuronal membrane was monitored with 2ms, 50pA test current pulses at 1Hz in current-clamp mode. Once a giga-ohm seal was formed, large zapping currents (five 200Hz 2ms 5nA pulses) were applied to break the cell membrane inside the whole-cell recording electrode. We used the following criteria to assess successful membrane breakthroughs and the attainment of proper whole-cell recordings: an immediate drop in voltage to the neuronal resting membrane potential, an instant decrease in voltage responses to test current pulses, a reduction in 50Hz mains AC noise, the appearance of spontaneous synaptic potentials and the presence of full-size action potentials overshooting to +20mV or above in response to current injections or during fictive swimming. During membrane resealing, these features disappeared and the whole-cell recording was lost. During more stable recordings, positive current injections were used to drive extra Ca^2+^ into the recorded neuron. Current steps were applied at 1Hz with amplitudes and durations ranging from 100-600pA and 300-600ms for 1-3 minute injection periods, depending on recording stability.

### Loose-patch extracellular recordings

For loose-patch recordings, gentle suction was applied to the neuron without breaking the cell membrane and spiking was recorded through evoked fictive swimming. dINs and non-dINs were classified by differences in their extracellular spike shapes (Soffe, Roberts and Li, 2009); dINs have longer peak-to-trough durations and smaller trough amplitudes (normalized to total peak-to-trough amplitude). One target dIN in each preparation was then whole-cell recorded to load 2 mM Ca^2+^. Target dIN identities were confirmed by neurobiotin staining following established methods (Li *et* al., 2001).

### Fluorescence analysis

Ca^2+^ imaging videos were exported to Fiji (Schindelin *et al*., 2012). XY drift was corrected with the Correct 3D Drift plugin (Parslow, Cardona and Bryson-Richardson, 2014) and regions of interest (ROIs) were drawn around the patched neurons. A large void region outside of neural tissue was used to calculate background fluorescence. The background fluorescence was then subtracted from the neuron ROI’s mean area pixel intensity to obtain fluorescence (F) values. The baseline fluorescence (F_0_) was calculated as the 10^th^ percentile F value 5 seconds before membrane breakthrough. ΔF/F as a percentage was then calculated as:

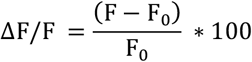

The following parameters were used to assess fluorescence dynamics after breakthrough: fluorescence increase was calculated as the maximum fluorescence after membrane breakthrough, rate of increase was the rate at which fluorescence reached its maximum value from breakthrough, and halftime was the time taken for fluorescence to drop to 50% maximum value.

### Pharmacology

In some experiments, we added 1 µM thapsigargin (Tocris Bioscience, Abingdon, UK), 10 µM SEA 0400 (Sigma-Aldrich, Gillingham, UK), 0.31 µM tetrodotoxin (TTX, Sigma-Aldrich, Gillingham, UK) and 150 µM Cd^2+^ (Sigma-Aldrich, Gillingham, UK) to the extracellular saline to block sarcoplasmic/endoplasmic reticulum Ca^2+^-ATPases (SERCAs) (Lytton, Westlin and Hanley, 1991), plasma membrane Na^+^/Ca^2+^ exchanger NCX1.1 (Bouchard *et al*., 2004; Lee *et al*., 2004), voltage-gated Na^+^ channels (Bane *et al*., 2014) and voltage-gated Ca^2+^ channels (Hinkle, Kinsella and Osterhoudt, 1987), respectively. This was done to facilitate Ca^2+^ diffusion to coupled cells and isolate loading-triggered Ca^2+^ increases from other physiological processes affecting cytoplasmic Ca^2+^. Thapsigargin inhibits all human SERCA family proteins (SERCA1, SERCA2a, SERCA2b, SERCA3) with equal potency (Lytton, Westlin and Hanley, 1991). The *Xenopus* SERCA2 has 92% percentage identity to human SERCA2 and the *Xenopus serca2/atp2a2* gene is expressed in neural tissue at stage 37/38 (Pegoraro, Pollet and Monsoro-Burq, 2011). SEA 0400 blocks NCX1.1, a human Na^+^/Ca^2+^ exchanger (Bouchard *et al*., 2004; Lee *et al*., 2004). We performed a BLAST analysis which revealed a *Xenopus* orthologue of NCX1.1, slc8a1.S, with high percentage identity (84.05%), which is expressed in tadpole neural tissue (Raciti *et al*., 2008).

### Statistical analysis

Shapiro-Wilk normality tests and Levene’s tests of equality of variances were used to select the appropriate statistical tests. For normally distributed data, Student’s *t*-tests were used. For non-normally distributed data, Mann-Whitney *U* tests and Kruskal-Wallis tests were used. Welch corrections for unequal variances and Holm corrections for multiple comparisons were used, when applicable. A Pearson’s correlation was used for regression analysis. All in-text descriptive statistics depict mean ± 1SD.

## Results

### Loading Ca^2+^ into neurons causes transient GCaMP6s fluorescence increases

We first loaded Ca^2+^ into random hindbrain and spinal neurons using whole-cell electrodes to see if this could trigger GCaMP6s fluorescence increases. We added 2 mM Ca^2+^ to the intracellular solution and performed whole-cell recordings with simultaneous Ca^2+^ imaging to capture fluorescence changes at the time of membrane breakthrough (Fig. 1A). We found significant increases in fluorescence upon breakthrough (148.6 ± 90.3%, *n* = 9) compared to standard intracellular solution with 0 mM free Ca^2+^ (14.4 ± 19.4%, *n* = 7, *p* < 0.01; Fig. 1B-C, Supplementary Movie S1). Next, to see if these dynamics differed to physiological single-cell activity, we compared fluorescence increases induced by Ca^2+^ loading and during fictive swimming (Fig. 1D). Fictive swimming bouts produced fluorescence peaks of smaller amplitude (Swim: 73.1 ± 9.3%, *n* = 9; Ca^2+^: 148.6 ± 90.3%, *n* = 9, *p* < 0.05; Fig. 1E) but faster rates of increase compared to Ca^2+^ loading (Swim: 62.7 ± 22.6 % s^-1^, *n* = 9; Ca^2+^: 19.5 ± 12.0 % s^-1^, *n* = 9, *p* < 0.001; Fig. 1F). Ca^2+^ loading-triggered fluorescence also had considerably longer halftime than fictive swimming (Swim: 2.7 ± 0.7 s, *n* = 9; Ca^2+^: 102.7 ± 84.1 s, *n* = 9, *p* < 0.01; Fig. 1G).

**Figure 1:**
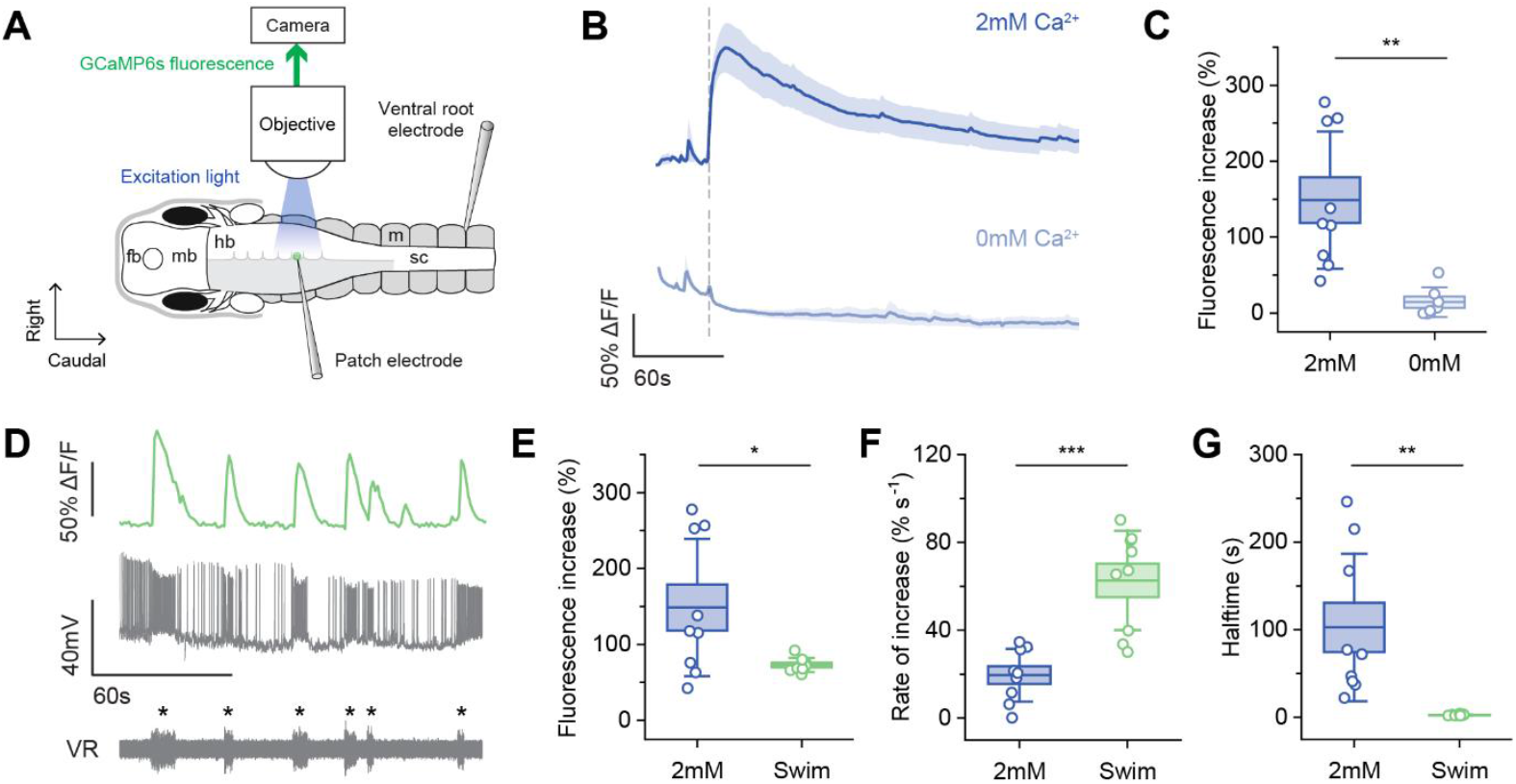
Loading Ca^2+^ into neurons through patch-clamp electrodes transiently increases GCaMP6s fluorescence. **A:** Dorsal view of a hemi-hindbrain tadpole CNS and diagram of the Ca^2+^ loading setup. The patch electrode is filled with intracellular solution containing 2 mM or 0 mM Ca^2+^. The ventral root electrode monitors fictive swimming activity. Fb: forebrain, mb: midbrain, hb: hindbrain, sc: spinal cord, m: myotome. **B:** Averaged (mean ± SEM) GCaMP6s fluorescence of the recorded neuron upon membrane breakthrough (vertical dashed line) with intracellular solution containing 2 mM and 0 mM Ca^2+^. 2 mM: *n* = 9, 0 mM: *n* = 7. **C:** Maximum fluorescence increase reached after 2 mM or 0 mM Ca^2+^ loading. Boxes: mean ± SEM, whiskers: ± SD; 2 mM: *n* = 9, 0 mM: *n* = 7; Student’s *t*-test with Welch corrections for unequal variances, ** *p* < 0.01. **D:** GCaMP6s fluorescence and whole-cell recording of a hindbrain neuron with simultaneous ventral root (VR) recording of six spontaneous fictive swimming episodes (marked with asterisks). **E-G:** Comparing fluorescence dynamics between Ca^2+^ loading and fictive swimming: maximum fluorescence increase, rate of increase and decay halftime. Boxes: mean ± SEM, whiskers: ± SD; 2mM: *n* = 9, Swim: *n* = 9; Student’s *t*-tests for normally distributed data (fluorescence increase, rate of increase), Mann-Whitney *U* test for non-normally distributed data (halftime), Holm corrections for multiple comparisons, * *p* < 0.05, ** *p* < 0.01, *** *p* < 0.001.

### Increasing intracellular Ca^2+^ concentration promotes membrane resealing

To maximize Ca^2+^ diffusion to potentially coupled neurons, we increased the Ca^2+^ concentration to 10mM. However, whole-cell recordings with 10 mM Ca^2+^ could not be sustained as signs of membrane resealing appeared which led to the loss of whole-cell recordings (see Materials and Methods; Fig. 2A). Upon re-examination, our 2 mM recordings also displayed indications of resealing (Fig. 2A). Whereas 100% of our 0 mM recordings could be held in whole-cell mode for over three minutes, 78% of 2 mM recordings showed resealing within three minutes and 100% of 10 mM recordings showed resealing within one minute of membrane breakthrough (Fig. 2B). Recordings with 10 mM Ca^2+^ were significantly shorter than those for 0 mM and 2 mM (Independent-samples Kruskal-Wallis test, *p* < 0.05 and < 0.001 respectively).

**Figure 2:**
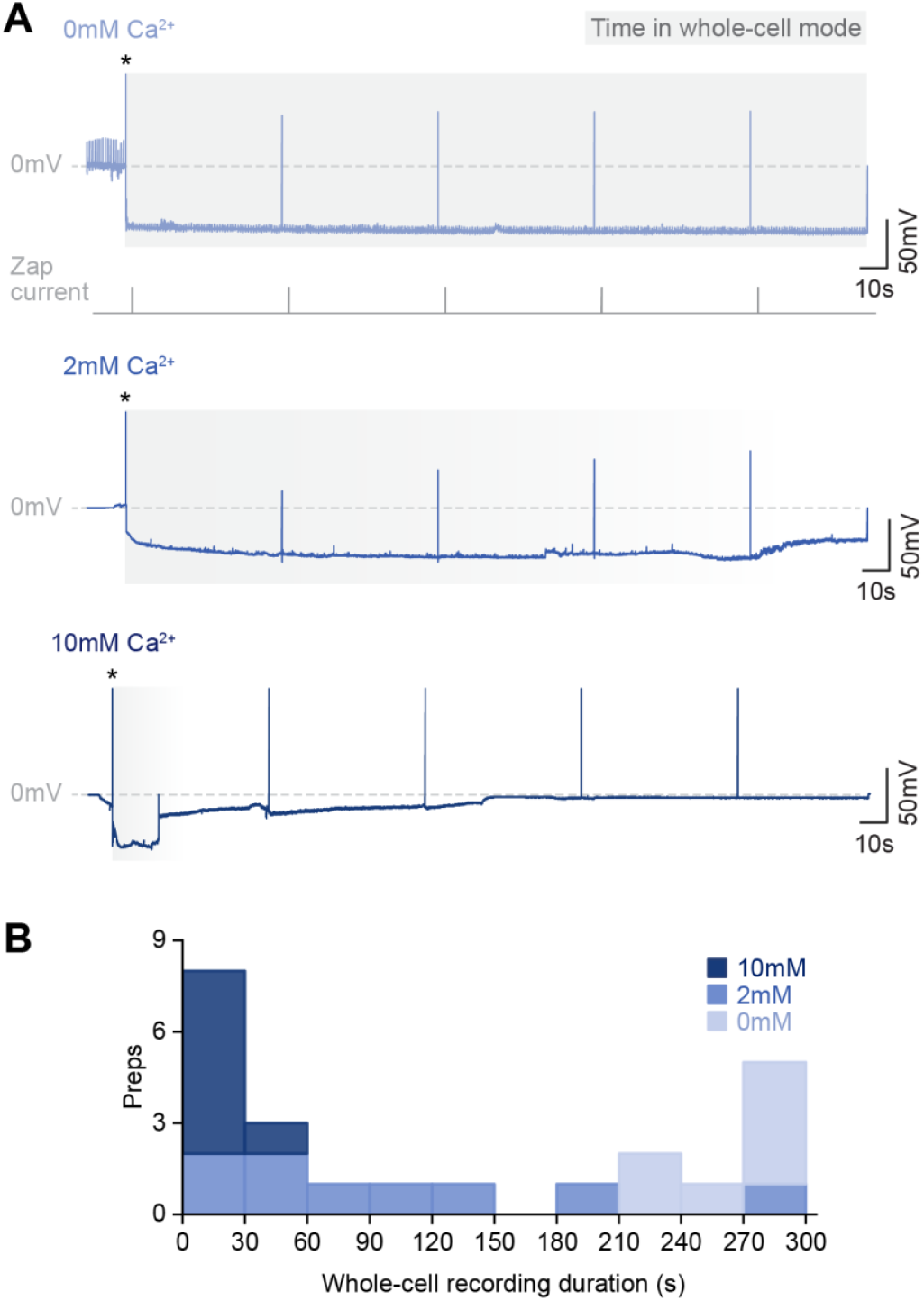
Increasing intracellular Ca^2+^ concentration promotes membrane resealing and loss of whole-cell recordings. **A:** Whole-cell recording traces with 0, 2 and 10 mM Ca^2+^ showing the progression of membrane resealing with increased Ca^2+^ concentration e.g., drift in membrane potential in 2 and 10 mM traces. Zap current is applied for membrane breakthrough to enter whole-cell mode and then to periodically monitor recording stability. **B:** Stacked histogram showing whole-cell recording durations for 0, 2 and 10 mM Ca^2+^. Duration starts from membrane breakthrough (marked by asterisks in A) and ends when physiological features characteristic of stable whole-cell recordings are lost (see Materials and Methods). 0 mM: *n* = 7, 2 mM: *n* = 9, 10 mM: *n* = 7.

### Blocking Ca^2+^ exchangers and pumps does not prolong fluorescence

Since stable whole-cell recordings could not be sustained with 10 mM Ca^2+^ intracellular solution, we applied blockers of Ca^2+^ exchangers and pumps to see if we could prolong fluorescence increases during 2 mM Ca^2+^ loading. Specifically, we used thapsigargin to block SERCA2s and SEA 0400 to block plasma membrane Na^+^/Ca^2+^ exchanger NCX1.1. In addition, we added TTX and Cd^2+^ to block voltage-gated Na^+^ and Ca^2+^ channels to remove activity-dependent Ca^2+^ influx and ensure fluorescence changes occurred only due to Ca^2+^ loading. Compared to our previous 2 mM Ca^2+^ loading experiments in physiological saline, the presence of blockers did not significantly affect maximum fluorescence increase (Saline: 148.6 ± 90.3%, *n* = 9; Drugs: 69.2 ± 28.7%, *n* = 8, *p* = 0.09; Fig. 3A-B) nor fluorescence decay halftime (Saline: 102.7 ± 84.1 s, *n* = 9; Drugs: 100.5 ± 80.8 s, *n* = 8, *p* = 1.0; Fig. 3C). Moreover, the duration of drug application had no effect on decay halftime (*r*^*2*^ = 0.20, *n* = 8, *p* = 0.15; Fig. 3D). Recordings were also similar in duration to those in control saline (Independent-samples Kruskal-Wallis test, *p* = 0.2, Fig. 3E).

**Figure 3:**
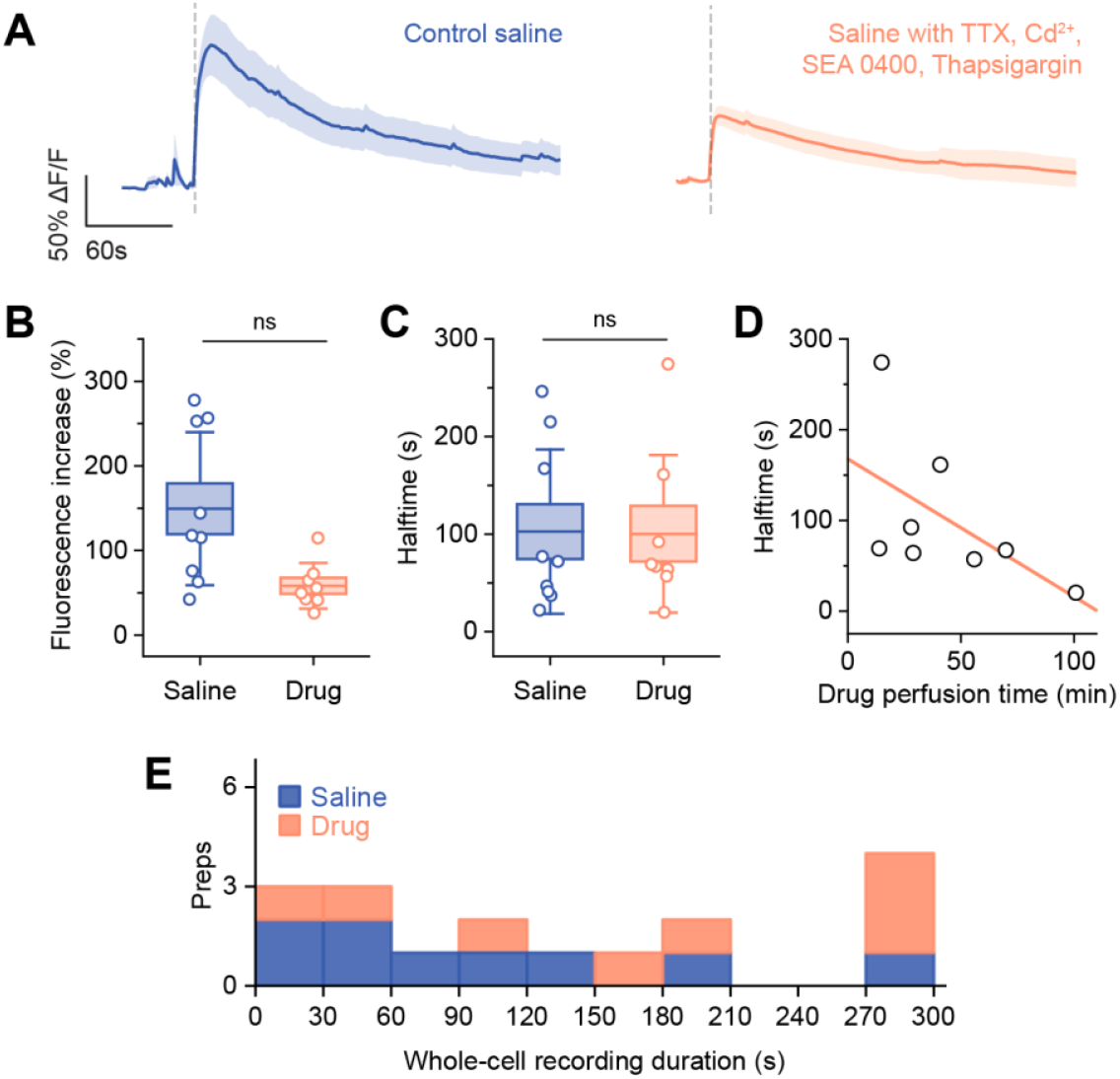
Ca^2+^ exchanger and pump blockers do not prolong fluorescence increases produced by 2 mM Ca^2+^ loading. **A:** Averaged (mean ± SEM) fluorescence upon membrane breakthrough (vertical dashed lines) in saline vs drug perfusion. **B-C:** Comparing maximum fluorescence increase and decay halftime in saline vs drug perfusion. Boxes: mean ± SEM, whiskers: ± SD; Saline: *n* = 9, Drugs: *n* = 8; Mann-Whitney *U* tests with Holm corrections for multiple comparisons. **D:** Scatter plot showing drug perfusion time before membrane breakthrough does not affect fluorescence halftime. Pearson’s correlation, *r*^*2*^ = 0.20, *p* = 0.15, *n* = 8. **E:** Stacked histogram showing whole-cell recording durations for saline vs drug perfusion. Duration starts from membrane breakthrough (marked by asterisks in A) and ends when physiological features characteristic of stable whole-cell recordings are lost (see Materials and Methods). Saline: *n* = 9, Drugs: *n* = 8.

### Ca^2+^ loaded into dINs does not spread to other dINs

Finally, we loaded Ca^2+^ into dINs and monitored if it could spread to neighboring dINs and increase their fluorescence. All neurons within the imaging field of view were first recorded in loose-patch mode to observe their spiking during swimming (Fig. 4B). Previous work found dIN spikes had a longer spike durations and smaller trough amplitudes than non-dINs (Soffe, Roberts and Li, 2009). In 27 neurons across 5 preparations, we identified 13 dINs and 14 non-dINs based on these criteria. Similar to previous findings, dIN peak-trough duration was longer (dINs: 3.3 ± 1.4 ms, *n =* 13; non-dINs: 0.8 ± 0.3 ms, *n =* 14, Mann-Whitney *U* test, *p <* 0.001) and trough amplitude was smaller (dINs: 0.3 ± 0.08, *n =* 13; non-dINs: 0.5 ± 0.07, *n =* 14, Mann-Whitney *U* test, *p <* 0.001) compared to non-dINs. Within each preparation, one target dIN was recorded in whole-cell mode and loaded with Ca^2+^, while other dINs and non-dINs (controls) in the field of view were monitored for fluorescence changes (Fig. 4A-C). dINs recorded in whole-cell mode were confirmed as dINs via morphological staining of their descending axons (Fig. 4D). We compared the fluorescence signals of the target dIN to surrounding loose-patched neurons (Fig. 4E). Target dINs showed increases in fluorescence after breakthrough (43.5 ± 24.0%, *n* = 5), which was significantly greater than surrounding dINs (8.0 ± 4.5%, *n* = 8, *p* < 0.01) and non-dINs (5.9 ± 3.8%, *n* = 14, *p* < 0.01; Fig. 4F). The fluorescence of these surrounding neurons did not differ from each other (*p* = 0.3) and remained unchanged even after three minutes (Fig. 4C).

**Figure 4:**
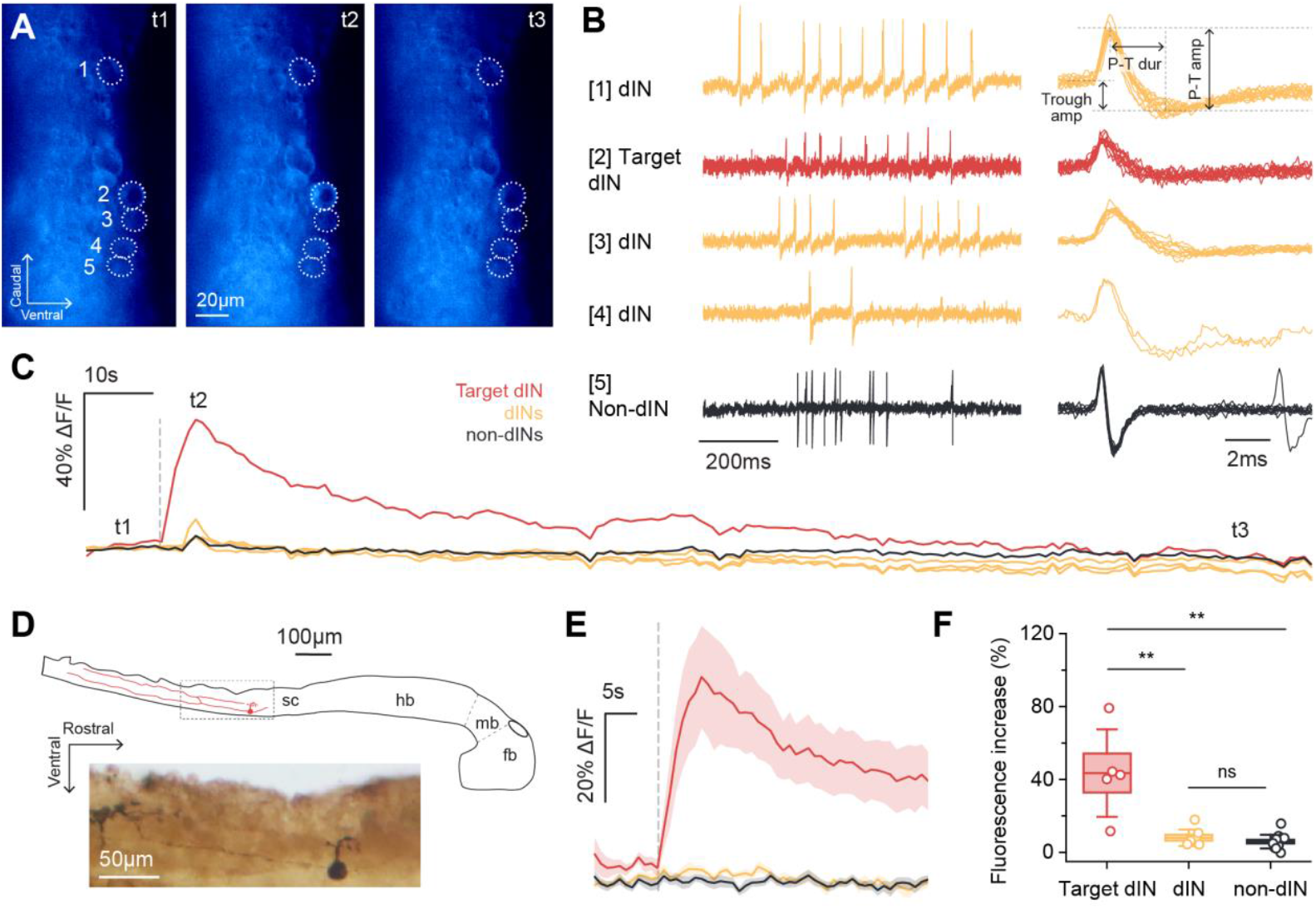
Ca^2+^ loading into target dINs does not lead to detectable fluorescence increases in other dINs and non-dINs. **A:** Fluorescence images of neurons (dashed circles) at timepoints t1-t3 in the fluorescence trace in **C**. Neurons 1-4 are dINs and Neuron 5 is a non-dIN based on extracellular spike shapes in **B**. Neuron 2 was recorded in whole-cell mode and loaded with 2 mM Ca^2+^. **B:** Left: loose-patch recordings for the 5 neurons in **A** during spontaneous swimming. Right: superimposed extracellular spikes for each neuron and spike shape parameters measured. P-T dur: peak-trough duration, Trough amp: trough amplitude, P-T amp: peak-trough amplitude. **C:** Fluorescence traces for the 5 neurons in **A** after membrane breakthrough in Neuron 2 over a three-minute period. Minimal fluorescence increases in other dINs and non-dINs at t2 are likely due to signal bleed-through from epifluorescence microscopy. **D:** Top: drawing of a whole-cell recorded dIN and its descending axon from neurobiotin morphological staining. Bottom: photograph of gray dashed box. Fb: forebrain, mb: midbrain, hb: hindbrain, sc: spinal cord. **E:** Averaged (mean ± SEM) fluorescence within one minute of membrane breakthrough for each neuron group. **F:** Comparing maximum fluorescence increases across neuron groups after target dIN breakthrough. Boxes: mean ± SEM, whiskers: ± SD; target dINs: *n* = 5, dINs: *n* = 8, non-dINs: *n* = 14; Kruskal-Wallis test with Mann-Whitney *U* post-hocs, ** *p* < 0.01.

In five different preparations, the whole-cell recordings were stable enough for us to inject positive current steps to drive extra Ca^2+^ into the dINs after the initial Ca^2+^ signal had faded. Current injections increased fluorescence in the patched dIN to its peak within a few seconds, followed by a plateau that lasted until the end of the current injection period (Fig. 5A-C). However, this did not produce fluorescence increases in neighboring neurons. The average fluorescence of target dINs across the current injection period (24.5 ± 15.0%, *n* = 5) was significantly higher than surrounding dINs (0.3 ± 2.5%, *n* = 9, *p* < 0.01) and non-dINs (1.1 ± 1.6%, *n* = 17, *p* < 0.001; Fig. 5D). As before, the fluorescence of surrounding dINs and non-dINs did not differ (*p* = 0.1; Fig. 5C).

**Figure 5:**
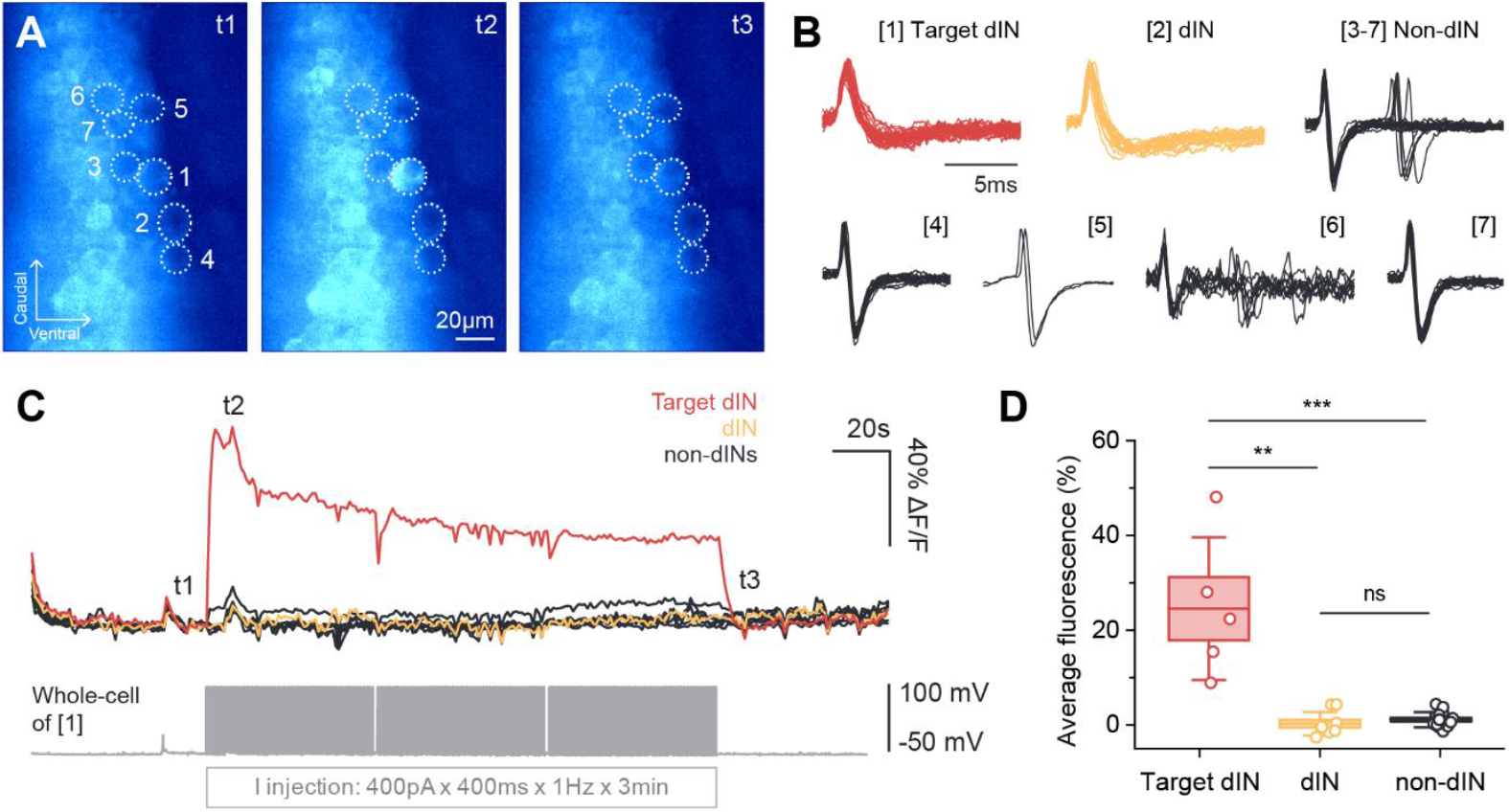
Extra Ca^2+^ loading through positive current injections into target dINs does not lead to detectable fluorescence increases in other dINs and non-dINs. **A:** Fluorescence images of neurons (dashed circles) at timepoints t1-t3 in the fluorescence trace in **C**. Neurons 1-2 are dINs and Neurons 3-7 are non-dINs based on extracellular spike shapes in **B**. Neuron 1 was recorded in whole-cell mode and loaded with 2 mM Ca^2+^. **B:** Superimposed extracellular spikes of the neurons in **A** during swimming. **C:** Fluorescence traces for the 7 neurons in **A** during three minutes of positive current injection into the loaded dIN (Neuron 1). **D:** Comparing average fluorescence during the current injection period across neuron groups. Boxes: mean ± SEM, whiskers: ± SD; target dINs: *n* = 5, dINs: *n* = 9, non-dINs: *n* = 17; Kruskal-Wallis test with Mann-Whitney *U* post-hocs, ** *p* < 0.01 *** *p* < 0.001.

## Discussion

In this study, we explored Ca^2+^ loading via whole-cell recording electrodes as a potential method for revealing electrically coupled neurons in GCaMP6s *Xenopus* tadpoles. First, we showed that loading 2 mM Ca^2+^ into neurons leads to an average fluorescence increase of 148.6% and an average decay halftime of 103 seconds. Ca^2+^ diffusion times across gap junctions have been reported previously. In coupled cochlear cells, it takes <1.5 seconds for elevated intracellular Ca^2+^ in one cell to lead to a detectable increase in Ca^2+^ in the other (Sun *et al*., 2005), whereas diffusion between coupled astrocytes and oligodendrocytes can take up to 155 seconds (Parys *et al*., 2010). Disparities between studies may reflect cell type, connexin composition, and methodological differences. In contrast to these studies investigating soma-somatically coupled cells, *Xenopus* dINs mostly have axo-axonal gap junctions (Li, Roberts and Soffe, 2009), which could be located hundreds of microns away from the soma and dendrites (Li and Soffe, 2019). More time and higher Ca^2+^ concentrations may be required for Ca^2+^ diffusion into the somas of coupled dINs.

We therefore increased the Ca^2+^ concentration in the intracellular solution to 10 mM. However, whole-cell recordings were extremely hard to maintain, and displayed signs of rapid membrane resealing after breakthrough. Resealing was also observed in our 2 mM recordings, although these could be held in whole-cell mode for longer. Ca^2+^ is required for many plasma membrane repair mechanisms after injury (Steinhardt, Bi and Alderton, 1994; Cheng *et al*., 2015). During membrane damage, an influx of extracellular Ca^2+^ (∼2 mM) into the cytosol (∼100 nM Ca^2+^ at rest) signals cell damage (Clapham, 2007). Ca^2+^ sensors like synaptotagmin VII, annexins and dysferlin detect rises in intracellular Ca^2+^ and promote various membrane repair mechanisms (Reddy, Caler and Andrews, 2001; Chakrabarti *et al*., 2003). Ca^2+^ chelators like EGTA and BAPTA can block membrane resealing by stabilizing free intracellular Ca^2+^, hence their inclusion in intracellular solutions for electrophysiology (Tsien, 1980). However, we excluded these from the intracellular solution to maintain high intracellular free Ca^2+^ and maximize diffusion. Consequently, Ca^2+^ loading likely triggered Ca^2+^-mediated membrane repair cascades, resulting in rapid loss of whole-cell recordings.

Instead of elevating Ca^2+^ concentration, we applied pharmacological blockers against *Xenopus* orthologues of SERCA2 proteins and NCX1.1. These blockers did not prolong fluorescence increase in the loaded cell, indicating the presence of active Ca^2+^ clearance mechanisms insensitive to these blockers. Many proteins help maintain Ca^2+^ homeostasis beside SERCAs and NCXs, including plasma membrane Ca^2+^ ATPases and Na^+^/Ca^2+^-K^+^ exchangers (Clapham, 2007). Adding more pharmacological blockers may sustain elevated cytoplasmic Ca^2+^ but may also pose greater risk to cell integrity. Instead, during stable recordings, we injected positive current steps to drive extra Ca^2+^ in the loaded cell. Although elevated fluorescence could be sustained in the target dIN throughout the injection period, no fluorescence increases were detected in neighboring cells, including dINs. This suggests that either more time is required for Ca^2+^ diffusion across axonal gap junctions or that Ca^2+^ diffused into coupled dINs was cleared too quickly or not detected by GCaMP6s. Indeed, detectable GCaMP6s fluorescence changes often requires bursts of action potentials (Fig. 1D) (Huang *et al*., 2021; Zhang *et al*., 2023). Single action potentials only are estimated to increase free cytoplasmic Ca^2+^ by 300-400 nM, lasting less than 2 ms (Helmchen, Borst and Sakmann, 1997; Sinha, Wu and Saggau, 1997). More sensitive genetically-encoded Ca^2+^ indicators may detect more transient, low concentration changes. This would, however, not overcome membrane resealing, which hinders introducing Ca^2+^ into the cell.

Gap junction permeability can also be regulated by cytoplasmic Ca^2+^. In many cases, high intracellular Ca^2+^ (nM to µM ranges) gates or uncouples gap junctions (Délèze and Loewenstein, 1976; Dahl and Isenberg, 1980; Peracchia, 2004). However, some studies on Cx36 and Cx43 gap junctions showed elevated intracellular Ca^2+^ is required to open channels and increase conductance (Pereda *et al*., 1998; Del Corsso *et al*., 2012; Wang *et al*., 2012). Which connexins form gap junctions between dINs remains unknown. Since swimming activity triggers increases in Ca^2+^ and electrical coupling strength between dINs does not change immediately after swimming (Li, Roberts and Soffe, 2009), it is unlikely that high intracellular Ca^2+^ decreases gap junctional conductances in dINs. Nevertheless, the rapid Ca^2+^ clearance and membrane repair mechanisms suggested here preclude Ca^2+^ loading as a technically feasible method for revealing gap junction connections. Novel genetic lines expressing genetically-encoded indicators of other ions (e.g., H^+^, K^+^, Cl^-^) (Kang and Baker, 2016; Shen *et al*., 2019; Lodovichi *et al*., 2022) with lesser impacts on cell homeostasis and integrity may be potential paths to explore. These have been used in previously mentioned “generator-reporter” methods (Kang and Baker, 2016; Wu *et al*., 2019), but are currently restricted to models susceptible to extensive genetic manipulation. Small, biologically inert molecules remain the most promising avenue. Many cell labeling studies have replaced organic dyes with quantum dots: inorganic fluorophores with broader excitation and narrower emission spectra (Jaiswal *et al*., 2003; Resch-Genger *et al*., 2008). However, these were also found to regulate Cx43 gap junctional communication, depending on delivery mechanism and cell culture setup (Chang, Hsu and Su, 2009). The need remains for more widely applicable and accessible methods.

To conclude, this study attempted to explore Ca^2+^ loading using whole-cell recording and Ca^2+^ imaging as a method for revealing electrically coupled *Xenopus* tadpole neurons. We encountered two fundamental challenges: 1) Ca^2+^ does not persist long in the loaded cell, likely due to rapid homeostatic clearance, and 2) loading Ca^2+^ induces loss of whole-cell recordings due to membrane resealing. As a result, fluorescence coupling could not be revealed between neurons previously shown to be electrically coupled. We therefore conclude that imaging Ca^2+^ diffusion is not technically viable for revealing coupled *Xenopus* tadpole dINs.

## Data Availability

Data can be found at the University of St Andrews Research Portal: https://doi.org/10.17630/4e0e1057-85f5-4793-8d95-78765fdde74b

## Supplemental Material

Supplementary Movie S1: https://youtu.be/Q8HU8QA8yUc

## Acknowledgements

We thank Prof Dave McLean and Dr Harmen Koning for their helpful discussions.

## Author Contributions

Conceptualization: Bella Xu Ying, Wen-Chang Li

Methodology: Bella Xu Ying, Wen-Chang Li

Investigation: Bella Xu Ying, Wen-Chang Li

Formal analysis: Bella Xu Ying, Wen-Chang Li

Visualization: Bella Xu Ying, Wen-Chang Li

Writing – original draft: Bella Xu Ying, Wen-Chang Li

Writing – review & editing: Bella Xu Ying, Wen-Chang Li, Maarten F. Zwart

Supervision: Wen-Chang Li, Maarten F. Zwart

